# Two-Year Field Testing of Genetically Engineered American Chestnut Reveals Fungal Blight Tolerance

**DOI:** 10.1101/2025.11.07.687213

**Authors:** Thomas Klak, Steven E Travis, Virginia Grace May, Ek Han Tan, Mathew WH Chatfield, Miranda Wheeler

## Abstract

The American chestnut (*Castanea dentata* [Marsh.] Borkh.) was a foundational forest canopy species in eastern North America until an accidentally imported fungal blight (caused by *Cryphonectria parasitica* [Murr.] Barr.) rendered it functionally extinct across its native range. Biotechnological approaches, such as the bioengineered Darling 54 line, have potential for future restoration of American chestnut, but field-based evaluations of blight tolerance have been limited. Progress has been slowed by the many years it takes for seedlings to grow to saplings, then to full-fledged trees. Current regulatory restrictions also constrain the testing of transgenic chestnuts to within permitted orchards. This research reports on a two-year field trial of Darling 54, their non-transgenic wild-type siblings and Chinese chestnut (*Castanea mollissima* Blume) controls deployed a randomized block design to test for blight tolerance. In the two years, three replicates each of 261 trees were branch-inoculated with EP-155, a highly virulent isolate of the fungal blight. Based on canker length, Darling 54 trees consistently outperformed their non-transgenic wild-type siblings and Chinese chestnut. To our knowledge, this is the first report of a multi-year field trial of fungal blight inoculations comparing advanced generation Darling 54 families. This field-based evaluation suggests that reintroduction programs using Darling 54 American chestnuts, which can commence after federal approval, may offer a promising path to success.

## Introduction

For millennia, the American chestnut (*Castanea dentata* [Marsh.] Borkh.*, Cd*) was a foundational species of eastern US forests stretching from Maine to Mississippi and west to Indiana (Russell 1987; Wang et al. 2013). In the late 1800s, a fungal blight that American chestnut is completely susceptiple to (caused by *Cryphonectria parasitica* [Murr.] Barr.) was accidentally introduced (Rigling and Prospero 2017). By the 1950s, the fungal blight had spread throughout the native range to kill the vast majority of the estimated four billion American chestnut trees (Powell et al. 2019). The species survives today mainly as stump sprouts, as the fungal blight does not kill the roots. However, trees rarely grow to maturity before succumbing to the blight (Dalgleish et al. 2016).

Genetic engineering is one strategy being developed to restore the American chestnut. Aiming to give American chestnuts enhanced fungal blight tolerance, researchers at SUNY-ESF^7^ inserted the Oxalate Oxidase (OxO) gene from wheat driven by various promoters via callus transformation into Ellis 1, a pure American chestnut line (Polin et al. 2006; Zhang et al. 2013; Newhouse et al. 2014). The expression of OxO appears to detoxify the oxalic acid that is excreted by *Cryphonectria parasitica* which is a necrotroph, thus conferring blight tolerance (Steiner et al. 2017). The transformation of Ellis 1, using a construct containing OxO driven by the strong, constitutive CaMV 35S promoter (Guilley et al. 1982) generated several transgenic events, including Darling 54 which is currently under consideration for federal deregulation (Jacobs et al. 2023; Newhouse et al. 2024; Klak et al. 2025). For this field trial, we utilized diversified trees derived from Darling 54 that were three or four breeding generations from the transformed T0 generation (Westbrook et al. 2020). We also included wild type full-siblings (null segregants) that did not inherit the Darling 54 transgene.

To date, no definitive fungal blight tolerance assays have been developed, but there are methods to approximate the impacts of natural fungal blight. Prior empirical work has determined that the optimal fungus inoculation methodologies vary significantly based on the age and stem diameter of the chestnut trees under study. Three of these inoculation assays, in order of the age and size of the tree for which they are best suited, are small stem, branch, and main stem. For young trees, a greenhouse based small stem assay has often been used, although this method has drawbacks. It has been found to have a large “no take” rate (i.e., inoculations that failed to transmit the fungus to the tree). It has also outright killed many tested seedlings and has tended to have a low correlation with later tests of larger trees (Conn et al. 2023; Newhouse et al. 2024). In fact, Steiner et al. (2017) showed that only T0 Darling 54 plants did not exhibit wilting or death after performing small stem assays, while the 50% of resistant ‘Qing’ variety of *Castanea mollisima* exhibited wilting or death. Second, for smaller trees, a branch inoculation assay known as the alternative small stem assay (AltSSA) can be utilized to measure the extent of fungal cankers and tissue injury as the distance it spreads down stems from the inoculation sites (Cipollini et al. 2021). It was developed and previously tested in Georgia, near the southern edge of the American chestnut’s range (Conn et al. 2023). The AltSSA yielded canker production at the site of inoculation of over 90% (i.e. few “no take” inoculations that are excluded from the data). This rate of successful inoculation is higher in many cases than other blight inoculation methods (Conn et al. 2023). Third, main stem inoculation has been used extensively for trees grown in the field with a 38mm (1.5in.) or larger diameter at breast height and is not recommended for smaller trees (Georgi and Fitzsimmons 2015; Penn State 2025).

In the present study, we utilized the AltSSA method because 69% of inoculated trees had stem diameters that are too small (<38mm) for main stem inoculation. For trees under 38mm diameter at breast height, which is the case for all the trees in this study, even highly resistant genotypes may be lethally girdled when inoculated at too small a diameter (Conn et al. 2023). Blight can aggressively penetrate through a small tree’s living layers rather than surrounding them as is typical fungal blight behavior with larger trees (Hebard 2012). The present study is also the first to deploy the AltSSA outside of the south, near the northern edge of the chestnut’s native range. Besides the AltSSA’s appropriateness for plants of sapling size, it has another advantage discovered in prior studies. Previous publications on the AltSSA describe it as “intended primarily for forward selection” which means that young chestnuts can be screened with the AltSSA and only those with small cankers can be outplanted, thus saving orchard space and effort. At the same time, the AltSSA deployed in previous work correlated with other measures of blight tolerance, including a multidimensional method gauging the blight tolerance of the young chestnut’s mother tree; the AltSSA also differentiated susceptibility between trees ranging from American chestnut to Chinese chestnut (*Castanea mollissima* Blume) and hybrids in between (Cipollini et al. 2021; Conn et al. 2023). To our knowledge, this is the first application of the AltSSA to compare genetically engineered chestnuts to full-sibling wild types.

To summarize, the AltSSA is deployed in the present research as the best nondestructive tool available to assess trees beyond seedling-size but not yet large enough for main stem inoculation in a field setting. The two main objectives of this study are: (1) to compare the fungal blight tolerance between Darling 54, non-transgenic wild-type American chestnut, and Chinese chestnut trees grown in a field plot in Maine, and (2) to evaluate the use of the AltSSA, a branch inoculation assay for quantifying the blight tolerance of American chestnut towards the ongoing efforts to breed blight tolerant trees for restoration.

## Materials and Methods

### Transgenic American Chestnut and Chinese Chestnut Trees Tested

The two-year field trial was conducted on a one-acre, APHIS^8^-permitted orchard (permit #124-833JJQQ) in long-term partnership with a private landowner in Cape Elizabeth, Maine (Klak 2018; TACF 2025; **Supp. Figs. 1 and 2** for general location). Chestnuts in this orchard came from wild type mother trees that had been bred the previous summer with Darling 54 pollen produced in labs at the University of New England and SUNY-ESF (Klak et al. 2025). The resulting seedlings were grown from nuts assembled from crosses with wild type trees in Maine and elsewhere provided by collaborators in 2021. A full-sibling family is defined here as a group of transgenic and non-trangenic (wild-type) progeny from a cross between a single pistillate tree and a single staminate hemizygous transgenic tree in the same season. Not all seedlings in the trial orchard survived the first or even the second year of outplanting and required replacement plantings of the following summers.

Tree height was measured to the nearest half centimeter from the top of the soil at the base of the tree to the woody tip of the tallest branch, which was usually the main stem. Stem diameter was measured with an electronic caliper to the nearest tenth millimeter approximately 16cm from the top of the soil. Because chestnut stems are often not exactly round, two measures were taken perpendicular to each other and averaged together.

One methodological improvement for Year 2 of the trial was the inclusion of additional surviving 4- to 5-year-old Darling 54 and wild-type orchard trees from the same families. This allowed for more intra-family comparisons of Darling 54 and wild-type siblings. At the same time, some trees that were challenged in Year 1 but did not have a full-sibling counterpart were excluded in Year 2. Lastly, all surviving trees possessing some proportion of Chinese chestnut alleles in the orchard were inoculated in Year 2. This increased the number of plants with Chinese chestnut alleles in the study from 19 in Year 1 to 30 in Year 2.

### Branch Inoculation Method based on the Alternative Small Stem Assay (AltSSA)

For this study, we deployed the EP-155 isolate of *Cryphonectria parasitica* (Crouch et al. 2020). It is the most virulent documented isolate of the chestnut blight and in recent years has been widely used in American chestnut resistance testing assays including the AltSSA (Hebard 2012; Conn et al. 2023). We performed branch inoculation using the AltSSA method that Cipollini et al. (2021) described, with minor changes noted here. EP-155 spores were obtained from West Virginia University’s Forest Pathology and Environmental Microbiology lab and were reconstituted in potato dextrose agar in petri dishes, following standard protocol (Kheyrodin 2024). The oven-plating procedure from Midwest Grow Kits (940 Dieckman St, Woodstock, IL) was used to plate spores onto agar in an aseptic environment. In the field, 4mm inoculum discs were cut to fit within the diameter of 5 or 6mm crimped straws and positioned so that the fungus was in direct contact with each newly cut branch (**Supp. Fig. 3**). In both years of the trial, we selected branches that were, whenever possible, south-oriented and mid-level on the tree. Branch diameters needed to be in the 4.5–5.5mm range to firmly secure the crimped straw containing the fungus plug against the fresh wound. Due to limited availability of branches, as well as internal checks of known resistant and susceptible trees (Chinese chestnut, wild-type siblings within Darling 54 families), we only conducted the AltSSA with the presence of EP-155.

### Canker Scoring Methods

In both years, trees were inoculated on June 5, and the crimped straws containing the inoculum were removed one week later as in previous studies (**Supp. Fig. 3**). After 90 days (September 5), inoculated stems were removed, and cankers and injured tissue were measured. The sites from where the stems were cut were disinfected to reduce the likelihood of further infection to the tree. Following protocol, two measurements were taken. One was the discolored canker zone where the fungus had advanced from the inoculation site down the branch. A second measure called full canker length was taken of a longer canker if there was evidence of additional, patchy discoloration up the branch as described by Cipollini et al. (2021) and Conn et al. (2023). The presence of sporulation on cankers was also recorded when orange clumps of spores, forming over the branches’ normally white lenticels, were visible (**Supp. Fig. 4**). If spores were present on one of the AltSSA inoculations, the entire individual was recorded for the presence of sporulation.

### Statistical Methods

Data were analyzed using R and followed a standard analysis of covariance (ANCOVA) approach, which treated chestnut type (i.e., Darling 54, wild-type siblings, or trees with Chinese chestnut alleles) as the principal factor of interest. Tree height was treated as a covariate of both of the canker length responses to fungal blight infection. In order to fully meet model assumptions for normality, Full Canker length measured in Year 1 and Height (both years) were square-root transformed, whereas Full Canker length measured in Year 2, Darkened Canker length and Orange Canker length were log-transformed.

ANCOVAs were first run using just the Darling 54 and wild-type families from both trial years to rule out family and year effects, and to thereby justify treating all trees used in our experiments as independent indicators of fungal blight tolerance. Although overall levels of replication were similar between our Year 1 and 2 datasets, the greater amount of replication *within* families conducted in Year 2 allowed for a comparison of chestnut types nested within their full-sibling families and year of planting. The lesser amount of within-family replication in Year 1 did not provide sufficient full-sibling families to allow for the same approach, but still allowed for separate comparisons of combined full- and half-sibling families by mother tree and by pollen source. The latter comparisons were made for Year 2 as well to assure consistency of results. These analyses indicated that neither year nor family was a significant variable in either the single Year 2 full-sibling model (**Supp. Table 1**) nor any of the Year 1 or 2 combined full- and half-sibling models (for mother tree results see **Supp. Table 2**; for pollen source results, see **Supp. Table 3**). These two factors, family and year, were therefore omitted from subsequent analyses, which allowed us to broaden our comparisons to include a third chestnut type, trees with Chinese chestnut alleles. Significant overall effects of chestnut type on discolored canker or full canker length were followed by multiple pairwise comparisons of type, which were evaluated by applying a Bonferroni correction to test-wise alpha levels. We sought to strike a balance between minimizing the probability of committing Type I and Type II errors and thus set our test-wise α-levels at 0.033 (α=0.10/3).

Prior to this study, genomic analysis (unpublished data obtained from genomics labs at the University of Notre Dame and Virginia Tech University) revealed that three of the wild type mothers of the orchard trees in the study had hybridity with European chestnut (*Castanea sativa* Mill.).^9^ Thirty seven of the sixty five (57%) trees tested in the Darling 54 families analysis in Year 2 of the trial had some European chestnut alleles, with the percentage ranging from approximately 10.5 to 20%. To determine if the European-American hybrids had a differing response to the blight than the 100% American chestnuts, a series of ANCOVAs was run, once again treating tree height as a covariate. Both chestnut type and percent European chestnut were run as uncrossed factors. Each blight response, as well as the tree height covariate, were log- or square-root transformed prior to analysis in order to fully meet model assumptions for normality.

## Results

### Tree height analysis

For both years, tree height and stem diameter were significant positive covariates with canker length on the branches that has been infected via the alternative small stem assay (AltSSA). In addition, tree height and stem diameter were highly correlated, as expected. We determined which of these two covariates had a more significant relationship with each response variable using a simple linear regression (response ∼ covariate). Tree height was more highly correlated with canker length than stem diameter so height was included in the model. Within family effects and the influence from the height covariate were factored out for all chestnut types, so the only difference between the Darling 54 and wild-type siblings was their presence or absence of the OxO transgene. When height was held constant and included in the model, Darling 54 trees had statistically shorter discolored cankers than wild-type trees (**Table 1**). The wild-type trees produced orange/darkened cankers that were on average ≥10mm (2024) and ≥6mm (2025) longer than their Darling 54 siblings (**Supp. Table 4)**.

**Table 1.** Full model and pairwise ANCOVA results from comparisons of all chestnut types measured for Orange/Darkened and Full Canker length in Year 1, and Darkened and Full Canker length in Year 2 based on type and tree height. Chestnut types include Darling 54, wild-type siblings (WT), and trees with Chinese alleles (CH).

| Comparison | Source | Year 1 Orange Canker |  |  | Year 1 Full Canker |  |  | Year 2 Darkened Canker |  |  | Year 2 Full Canker |  |  |
| --- | --- | --- | --- | --- | --- | --- | --- | --- | --- | --- | --- | --- | --- |
|  |  | F | DF | p | F | DF | p | F | DF | p | F | DF | p |
| All | Type | 17.58 | 2,161 | <0.0001*** | 22.10 | 2,161 | <0.0001*** | 8.85 | 2,92 | 0.0003*** | 7.19 | 2,92 | 0.0013** |
|  | Height | 5.99 | 1,161 | 0.0155* | 8.98 | 1,161 | 0.0032** | 8.07 | 1,92 | 0.0056** | 6.60 | 1,92 | 0.0118* |
| Darling 54/WT | Type | 32.90 | 1,143 | <0.0001*** | 39.55 | 1,143 | <0.0001*** | 5.61 | 1,63 | 0.0209* | 3.31 | 1,63 | 0.0737 |
|  | Height | 8.39 | 1,143 | 0.0044** | 11.69 | 1,143 | 0.0008*** | 10.86 | 1,63 | 0.0016** | 6.46 | 1,63 | 0.0135* |
| Darling 54/CH | Type | 2.12 | 1,110 | 0.1478 | 6.58 | 1,110 | 0.0117* | 16.80 | 1,61 | 0.0001*** | 13.81 | 1,61 | 0.0004*** |
|  | Height | 4.02 | 1,110 | 0.0473* | 6.06 | 1,110 | 0.0154* | 4.22 | 1,61 | 0.0443* | 6.05 | 1,61 | 0.0167* |
| WT/CH | Type | 8.34 | 1,68 | 0.0052** | 5.13 | 1,68 | 0.0266* | 3.84 | 1,59 | 0.0549 | 3.97 | 1,59 | 0.051 |
|  | Height | 0.35 | 1,68 | 0.5575 | 0.73 | 1,68 | 0.3964 | 3.14 | 1,59 | 0.0814 | 1.67 | 1,59 | 0.2008 |

The tree height-stem diameter correlation among all Darling 54 and wild-type trees was 0.833 (n=70). Chinese chestnuts tend to grow wider and shorter than American chestnuts. When trees with Chinese alleles were included (n=96), and the correlation was lower (0.785) but still high. Between height and diameter, height is worth examining more closely to see if the Darling 54 were shorter than their wild-type siblings, which has been reported elsewhere (TACF n.d.), and was attributed to costs of defending against pathogens that can consume a plant’s energy and resources that would otherwise be put to growth (Monson et al. 2022). In our experimental orchard, Darling 54 trees had a mean height of 195.3cm (sd=85.3cm) while their wild-type siblings had a similar mean height of 190.4cm (sd=70.3cm). The standard deviations indicate considerable height variation in each group, but Darling 54 trees were not on average shorter than their wild-type siblings. As has been recently reported in a study of chestnut lines variously planted in open field and shelterwood sites in three other US states, chestnut growth and performance will vary per location and site conditions (Olichney et al. 2026).

### Year 1 AltSSA Canker Measurement and Analysis

Canker lengths were measured from the inoculation site on the branch at two distances: orange canker and full canker (**Supp.Fig 4**). The most significant contrast observed was between the Darling 54 and wild-type siblings. They differed statistically for both measures of canker length (orange canker length: F1,143=32.90, p<0.0001; full canker length: F1,143=39.55, p<0.0001; **Table 1** and **Fig. 1**). Darling 54 also had significantly shorter full canker lengths than trees with Chinese alleles (F1,110=6.58, p=0.0117; **Table 1** and **Fig. 1**).

**Fig. 1.**
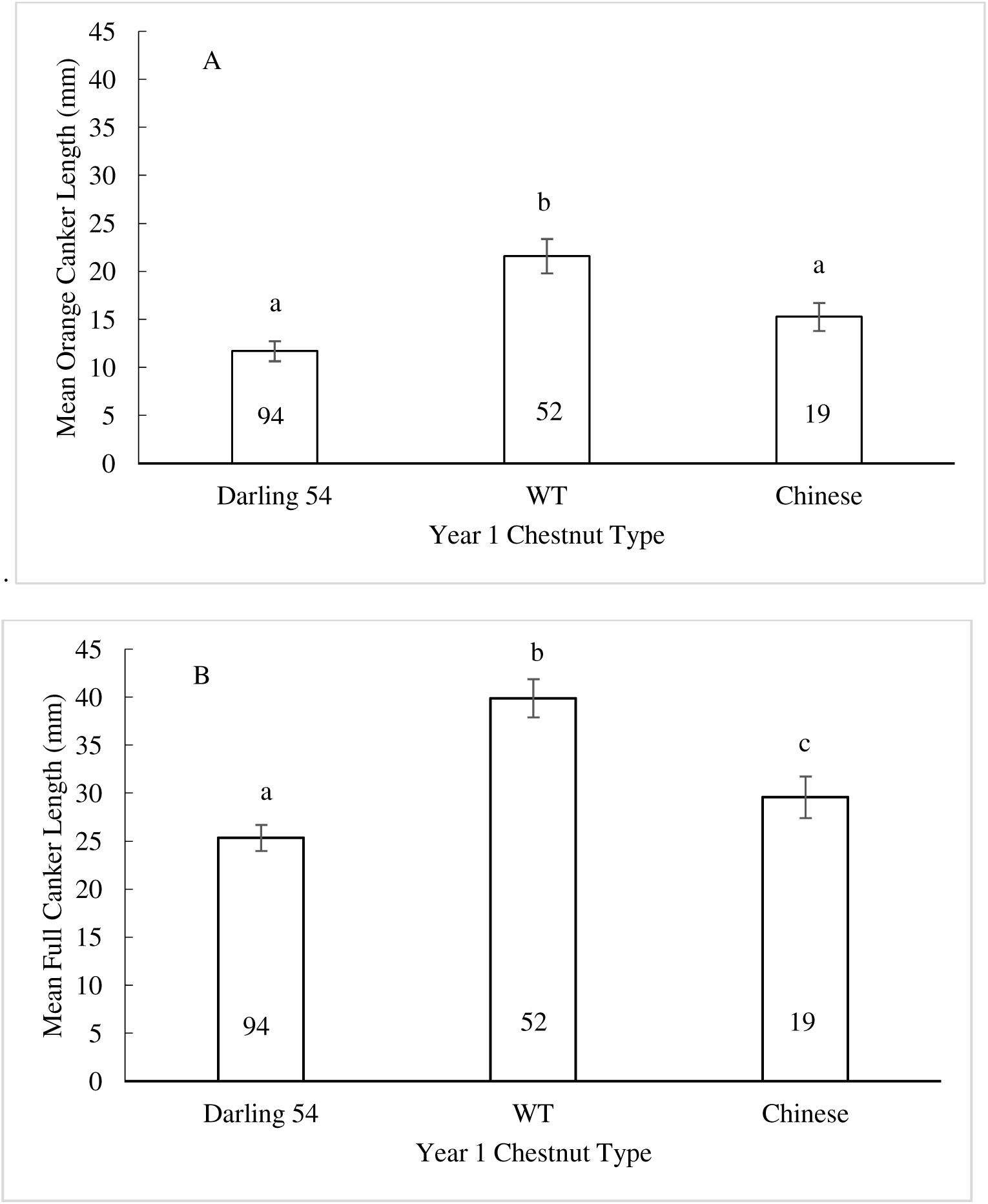
90-day mean canker lengths for the three chestnut types in Year 1. Fig1A: Orange canker. Fig1B: Full canker. The number inside each bar graph is the sample size. Darling 54 are hemizygous lines that carry the 35S::OxO transgene, WT refers to null segregants from the same families that are wild-type, and Chinese refers to plants with Chinese chestnut alleles. Error bars represent standard errors. Shared letters above the bars indicate non-statistically-significant differences in means.

The presence of sporulation on cankers was also recorded. If spores were present on one of a plant’s inoculations, the entire individual was recorded for the presence of sporulation. A χ2 test was run on the presence or absence of sporulation to compare chestnut types, then a series of odds ratio tests was run as planned contrasts to determine if Darling 54 trees differed in sporulation presence from their wild-type siblings or the chestnuts with Chinese alleles. The only significant result of the three planned contrasts was between Darling 54 and wild-type siblings. There was significantly less sporulation observed on Darling 54 trees compared to wild-type siblings (odds ratio 0.403, confidence interval 95% [0.181, 0.901], p=0.013).

### Year 2 AltSSA Canker Measurement and Analysis

We observed that canker discoloration from fungal blight infection manifested differently in the two years of the study. In Year 1, the zone appeared orange and aligned with coloration reported in other studies (May 2025). In Year 2, the discoloration was dark and the texture was more shriveled. We speculate that the more shriveled and darker-colored branches were impacted by the 2025 drought. Consistent with this interpretation is the fact that whereas fungal sporulation was observed on most (i.e., 69%) orange zone branches in Year 1, it was entirely absent in the Year 2 drought year. We observed that in Year 2 the fungal inoculations appeared phenotypically drier on the branches. To capture the inter-year differences, we analyzed the darkened canker for the Year 2 on branches inoculated via AltSSA.

Based on this analysis, the darkened cankers on Darling 54 trees in Year 2 were significantly shorter than those on wild-type siblings (F_1,63_=5.61, p=0.0209) and also significantly shorter than those of trees with Chinese chestnut alleles (F_1,61_=16.80, p=0.0001; **Table 1** and **Fig. 2**). Whether the canker zone was orange as in Year 1 or darkened as in Year 2, there is considerable consistency in the inter-year findings when comparing Darling 54 trees to their wild type siblings (**Figs. 1** and **2**). As noted, fungal sporulation differed between Years 1 and 2. Sporulation was observed on most (i.e., 69%) orange zone branches in 2024, it was entirely absent at 90 days in 2025. Like the chestnut hosts, the fungal blight appeared phenotypically drier during the 2025 drought.

**Fig. 2.**
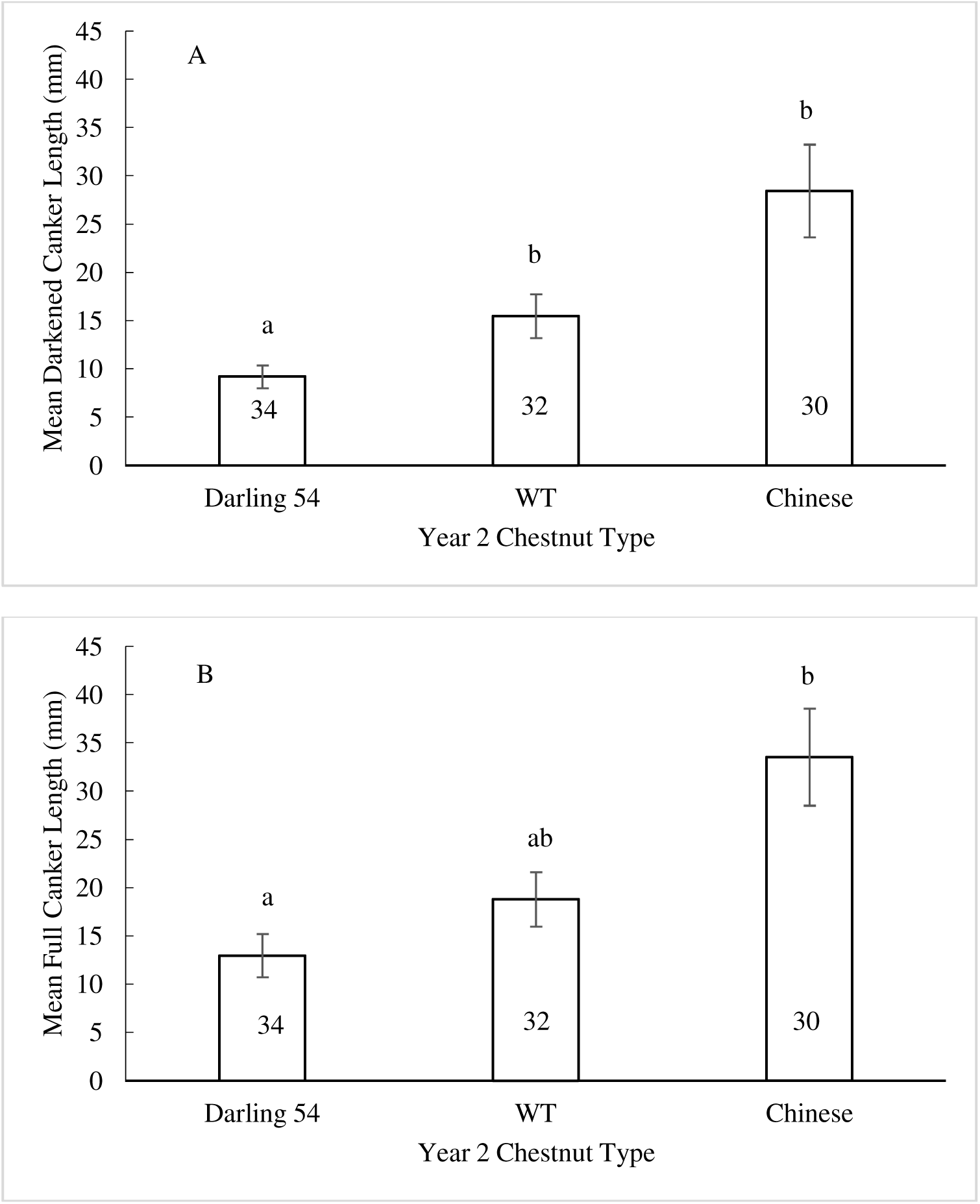
90-day mean canker lengths for the 3 chestnut types in Year 2. Fig2A: Darkened canker. Fig2B: Full canker. The number inside each bar graph is the sample size. Darling 54 are hemizygous lines that carry the 35S::OxO transgene, WT refers to null segregants from the same families that are wild-type, and Chinese refers to plants with Chinese chestnut alleles. Error bars represent standard errors. Shared letters above the bars indicate non-statistically-significant differences in means.

Another question this study addressed was whether cankers differed between trees that were 4- versus 5-years-old in Year 2. A well-replicated test of outplanting year and family via comparison of Darling 54 and wild-type siblings showed no effect of either factor on blight canker metrics (e.g., for darkened canker length, Year: F_1,10_=0.06, p=0.8064; Family: F_10,12_=0.31, p=0.9646; **Supp. Table 1**). After these alternative explanations were discounted, the conclusion was that both canker measurements showed that Darling 54 trees outperformed their wild-type siblings (F_12,40_=2.69, p=0.0095; **Supp. Table 1**).

### Influence from European chestnut on AltSSA canker length

As mentioned earlier, genomic analysis revealed that three of the wild type mothers of the orchard trees in the study had hybridity with European chestnut (*Castanea sativa* Mill.). Thirty seven of the sixty five (57%) trees tested in the Darling 54 families analysis in Year 2 of the trial had some European chestnut alleles, with the percentage ranging from approximately 10.5 to 20%. European chestnut is also thought to be susceptible to *Cp* and should therefore not contribute resistance alleles in these trees (Ježić et al. 2024). We therefore investigated the question of whether the percentage European chestnut genes could explain fungal AltSSA canker length. A series of ANCOVAs was run, once again treating tree height as a covariate. In both years, canker lengths from trees that inherit European chestnut alleles was not statistically significant compared to canker lengths of 100% American chestnut trees (with p-values ranging from 0.30–0.47; **Supp. Table 4**).

## Discussion

In this study, we showed that Darling 54 outperformed wild-type American chestnut and Chinese chestnut over two years of field trial based on the alternative small stem assay (AltSSA), using the most virulent *Cryphonectria parasitica* isolate, EP-155. Over the two years, we measured canker lengths on 261 trees in triplicates from AltSSA infections and showed that Darling 54 exhibited significantly smaller canker lengths compared to checks. Previous field tests of Darling 54 on fungal blight tolerance are few and based on small-scale field tests (e.g., only n=3 Darling 54 trees and n=3 wild-type siblings) that were conducted on first or second generation Darling 54 trees in a single year (Powell et al. 2020; ESF 2023; Newhouse et al. 2024).

Methodologically, this study provides further support for the efficacy of Cipollini et al.’s (2021) branch inoculation assay (AltSSA), specifically as a tool for assessing chestnut trees in an orchard setting. The successful inoculate rate of the AltSSA is high. The two previous AltSSA studies had very low “no take” rates and ≥98% inoculation success (Cipollini et al. 2021; Conn et al. 2023). In our study, the “no take” rates were slightly higher, with a 91% inoculation success rate in Year 1 and 95% in Year 2, which is comparable to these studies. In comparison, our AltSSA “no take” rates were much lower than the ∼40% rate produced by traditional small stem assays (Gentner 2018). This study demonstrated the ability of the AltSSA to yield meaningful results in a different outdoor environment at the northern edge of the species’ range. Furthermore, we were able to improve on previous studies by inoculating each sapling three times, with each being treated as a subsample. The subsampled canker measurements were averaged together to reduce variation and more inclusively detect blight response. Lastly, the AltSSA has previously been used to compare blight resistance among American and Chinese chestnuts and their hybrids. This study was the first to establish a baseline response for blight tolerance of advanced generation Darling 54 trees compared to full-sibling wild-type trees in the field.

Our results from both years showed that Darling 54 trees outperformed trees with Chinese alleles when using the AltSSA method and EP-155, although this finding requires further investigation because of the small number of trees with Chinese alleles used in this comparison. In three of the four comparisons in the two years, Darling 54 significantly outperformed trees with Chinese chestnut alleles (**Table 1** and **Figs. 1-4**). This might be viewed as an unexpected finding, considering that the Chinese chestnut coevolved with the fungal blight and therefore should have natural blight tolerance. However, some previous studies using the small stem assay found a similar pattern, including the SUNY-ESF submission to APHIS for nonregulated status for Darling 54 (Newhouse et al. 2024). The American Chestnut Foundation also reported that Darling 54 (erroneously referred to then as Darling 58) had shorter cankers than full Chinese chestnuts, although the difference was not statistically significant in their small study (TACF 2023). The comparison of Darling 54 and trees with Chinese chestnut alleles is a subject especially in need of further testing.

A major complication for assessing fungal blight tolerance is the American chestnut’s natural lifespan of five hundred or more years in the absence of blight. Predictions about long-term survivability based on evidence from young trees must be taken with caution. Nonetheless, it is valuable for making incremental progress toward the goal of chestnut restoration to assess the fungal blight tolerance of younger trees. This is particularly true in the field where, instead of the relatively stable, consistent, mild and idealized lab/greenhouse conditions, trees are exposed to harsher weather, seasonality, herbivory, a wider range of environmental conditions, and vicissitudes such as late freeze and summer drought. The young trees evaluated in this study have already experienced all of the above.

Unfavorable weather conditions significantly impacted younger chestnut trees across Maine in 2025 (Year 2). Whereas 2024 (Year 1) was a relatively normal summer for temperatures and rainfall by historical standards, 2025 was exceptionally difficult for younger chestnuts. Compared to 2024, 2025 had greater swings from a wet early summer followed by prolonged drought. Commenting on the impacts on agriculture in 2025, which would also apply to young chestnut trees, Maine’s Drought Task Force explained that “Crops typically require about 1” of water per week and this has not happened during June through August” (Maine 2025; see also **Supp Figs. 1** and **2**). The timing of the drought corresponded with the field trial in Year 2 and these impacts on chestnut trees of the extreme weather were visible during the 90-day period in Year 2. Wilted leaves, leaf drop, seedling death, and, for mature trees outside the orchard, bur drop and the aborting of flowers were observed. Across five monitored WT orchards that host 5- to 8-year-old trees in different Maine locations, the production of burs was severely diminished and even zero in some places (data not shown). In prior, more normal years, there was an increasing crop of flowers followed by burs and fertile nuts on trees as they matured, as expected with chestnut maturation. These observations, along with what we observed in terms of disease development on these trees in 2025 compared to 2024, are potentially associated with drought, resulting in an atypical growing season in 2025.

In conclusion, we suggest that experimentally challenging Darling 54 with highly virulent fungal isolates such as EP-155 should extend from the baseline established in this two year trial. Additional trials that can provide further documentation of how Darling 54 trees perform under varying site conditions is warranted. One recommended next step is to continue to challenge Darling 54 trees as they grow older and larger. Another is to challenge trees growing in different geographical locations across the American chestnut native range. A third is to evaluate how differing site specific environmental conditions, such as soils and varying amounts of light and shade, impact Darling 54 performance. Indeed, other recent empirical work focused on tree growth and survival has documented that geographical and site conditions shape the relative performance of various chestnut lines including Darling 54 (Olichney et al. 2026). As the American chestnut restoration project continues to move incrementally forward, additional trials will provide further insights as to the extent to which the OxO gene offers sufficient fungal blight protection to Darling 54 American chestnuts.

## Author Contributions Statement

T.K., V.M., E.H.T., M.C. and M.W. conducted the field trial. T.K., S.T. and V.M. prepared the data. S.T. and V.M. analyzed the data. T.K. and S.T. prepared the tables and figures. T.K., E.H.T, and V.M. wrote the manuscript. All authors reviewed and edited the manuscript.

## Acknowledgments

We thank colleagues at SUNY-ESF (State University of New York – Environmental Science and Forestry) for providing Darling 54 seeds and seedlings, and at West Virginia University’s Forest Pathology and Environmental Microbiology lab for providing the chestnut blight fungus. For helpful suggestions on earlier drafts, the authors thank Jack Kennell, Jay Wason III, and three anonymous reviewers.

## Supplemental Material

**Supplemental Fig. 1.**
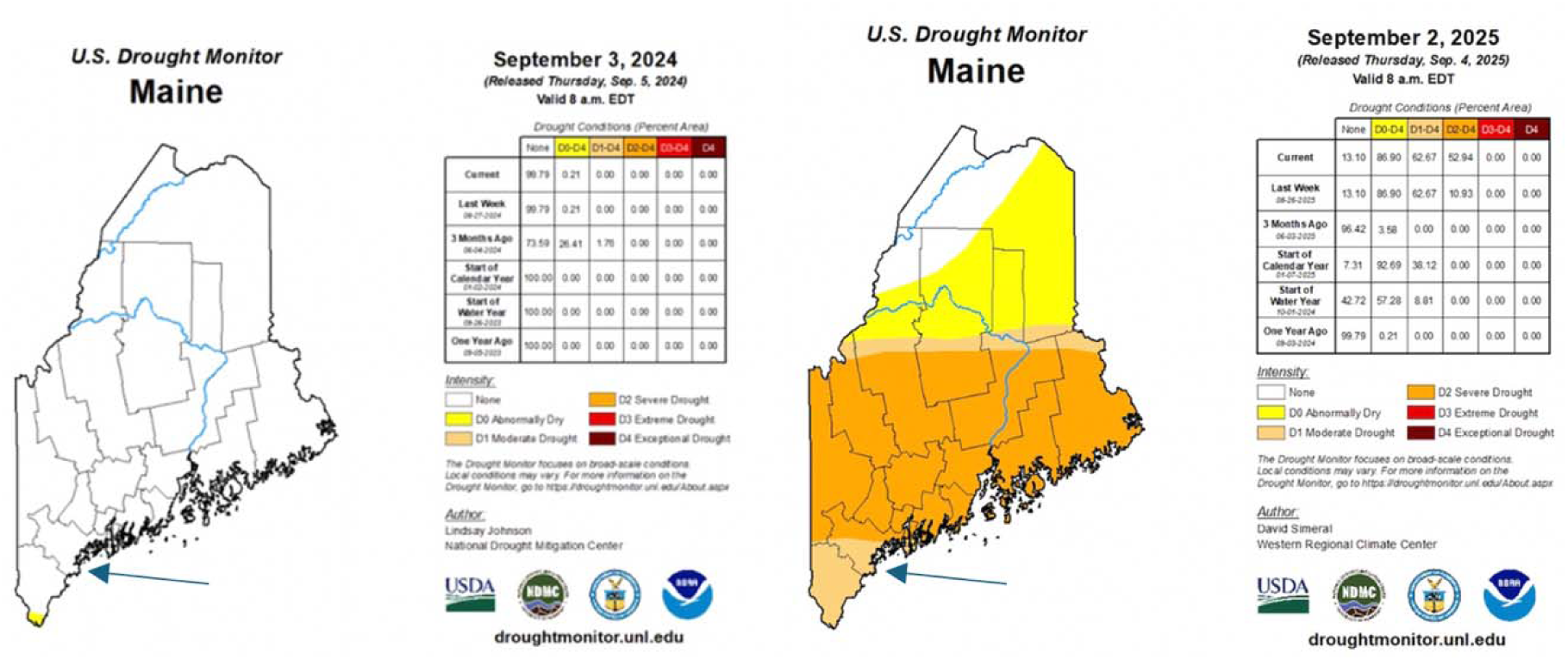
Moisture Conditions in 2024 and 2025 by the end of the 90-day inoculation period. Arrow points to the field trial orchard’s location Source: US Drought Monitor

**Supplemental Fig. 2.**
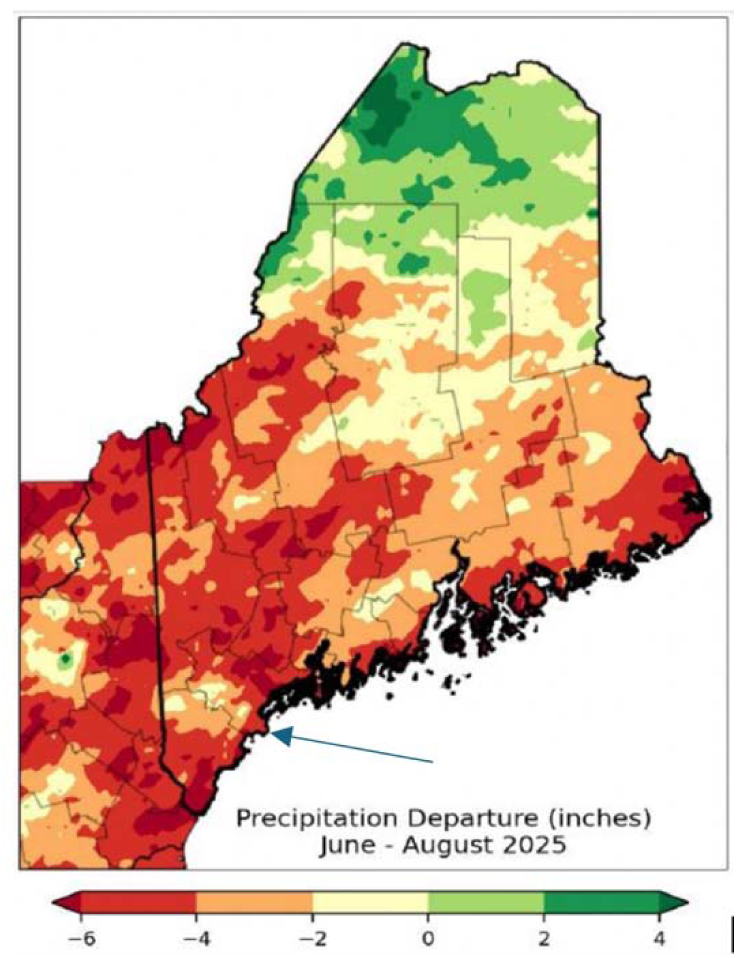
Precipitation deficit during the 2025 90-day inoculation test. Arrow points to the field trial orchard’s location Base Map Source: Maine, 2025

**Supplemental Fig. 3.**
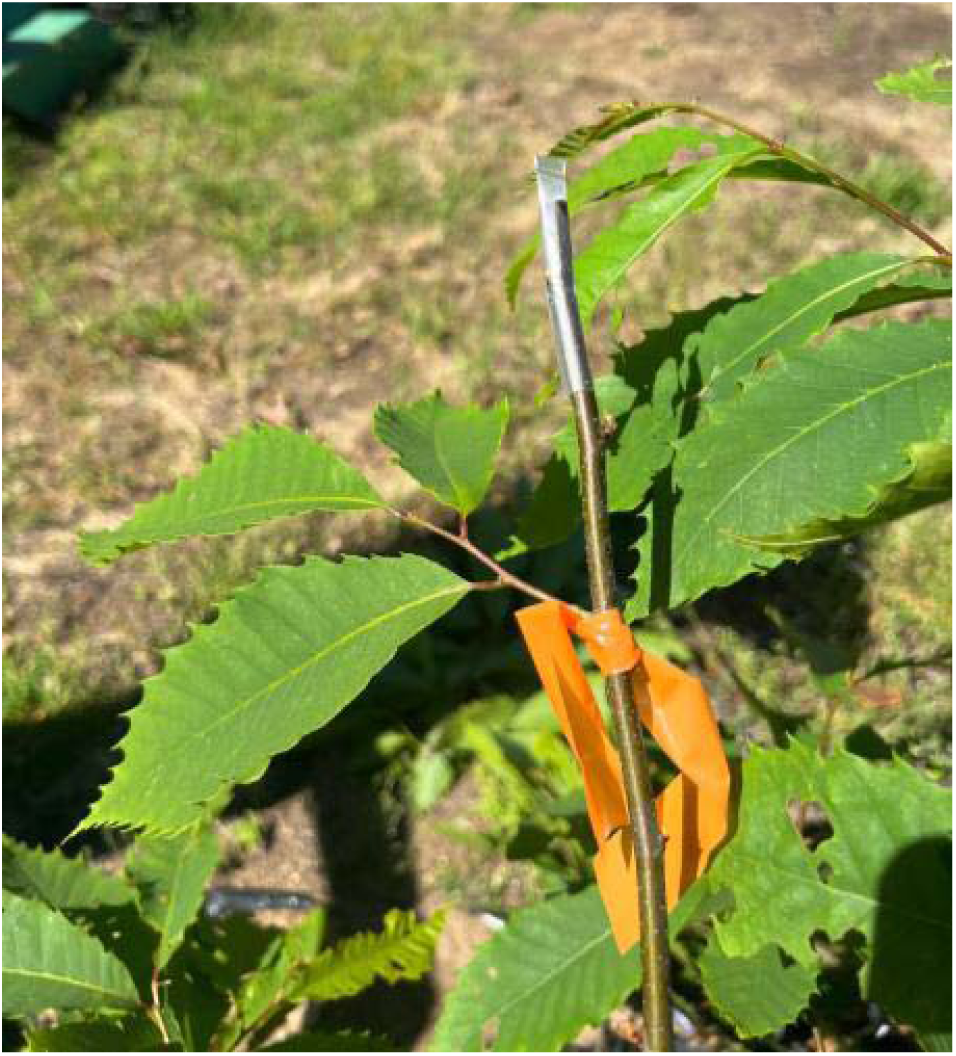
Flagged branch immediately after inoculation, with crimped, clear plastic straw sealing the inoculum disc snugly to the cut end of branch

**Supplemental Fig. 4.**
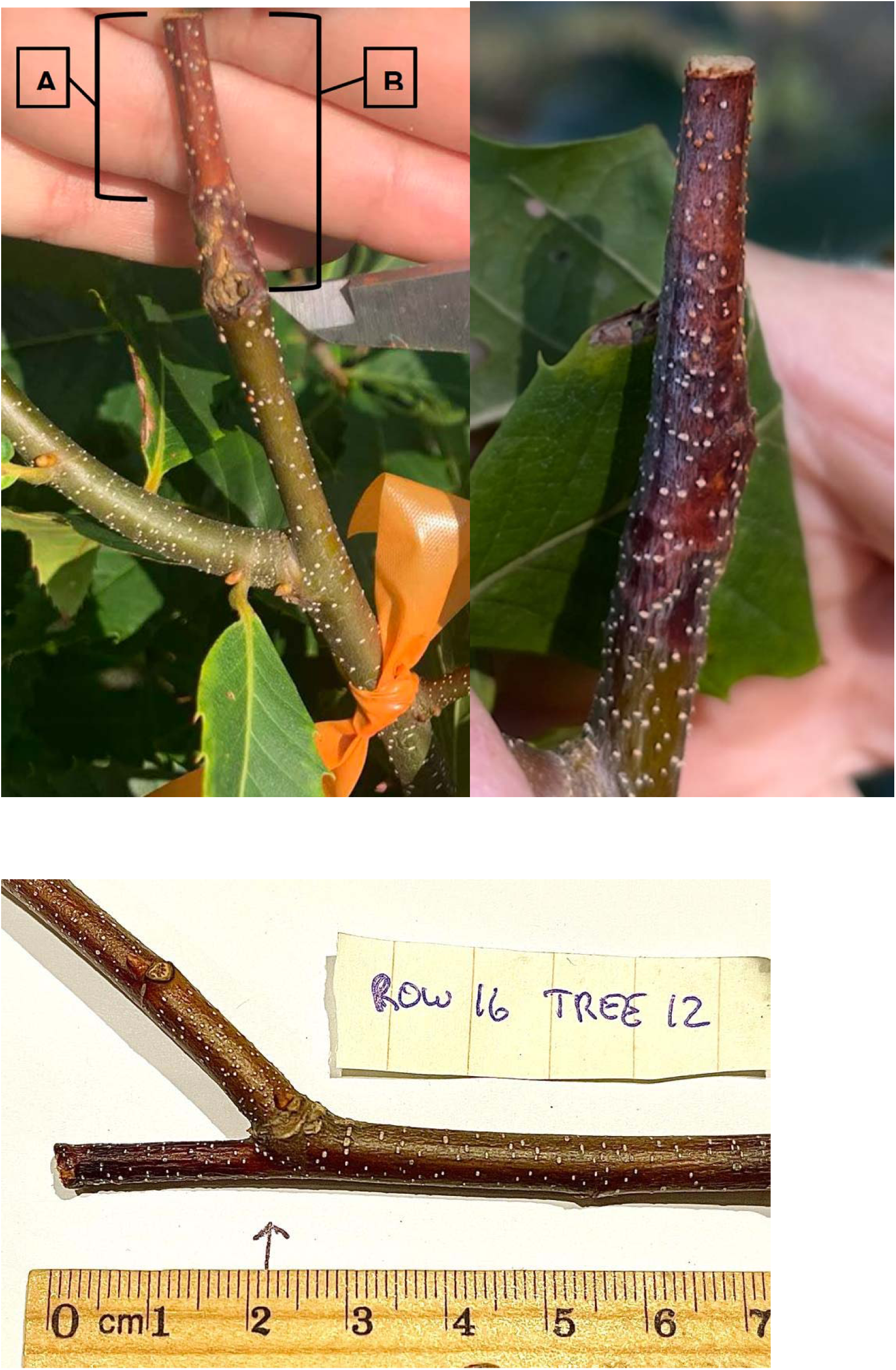
Example cankers from 2024 (top; A=orange canker length including normally-white lenticels turned orange indicating sporulation in the right photo; B=full canker length); and 2025 (bottom) showing darkened, more shriveled canker where in this case both canker lengths are nearly the same

## Supplemental Tables

**Supp.Table 1.**
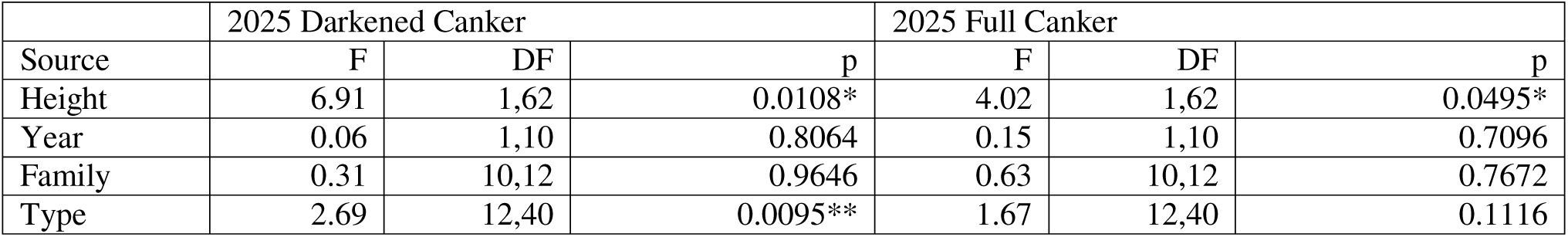
Full-sib Nested Analysis of Covariance (ANCOVA) results evaluating the effects of Year, Family within Year, and Chestnut Type within Family (including Darling 54 and Wild-Type) on Darkened and Full Canker length measured in Year 2. Due to the nested nature of the factors, the ANCOVA’s were run in stepwise fashion, first evaluating the effect of the covariate, height, on each branch inoculation assay via Simple Linear Regression analysis, and then running Nested ANOVAs on the residuals from the regressions. In order to fully meet model assumptions, Height was square-root transformed, whereas Darkened and Full Canker length were log-transformed.

|  | 2025 Darkened Canker |  |  | 2025 Full Canker |  |  |
| --- | --- | --- | --- | --- | --- | --- |
| Source | F | DF | p | F | DF | p |
| Height | 6.91 | 1,62 | 0.0108* | 4.02 | 1,62 | 0.0495* |
| Year | 0.06 | 1,10 | 0.8064 | 0.15 | 1,10 | 0.7096 |
| Family | 0.31 | 10,12 | 0.9646 | 0.63 | 10,12 | 0.7672 |
| Type | 2.69 | 12,40 | 0.0095** | 1.67 | 12,40 | 0.1116 |

**Supp.Table 2.**
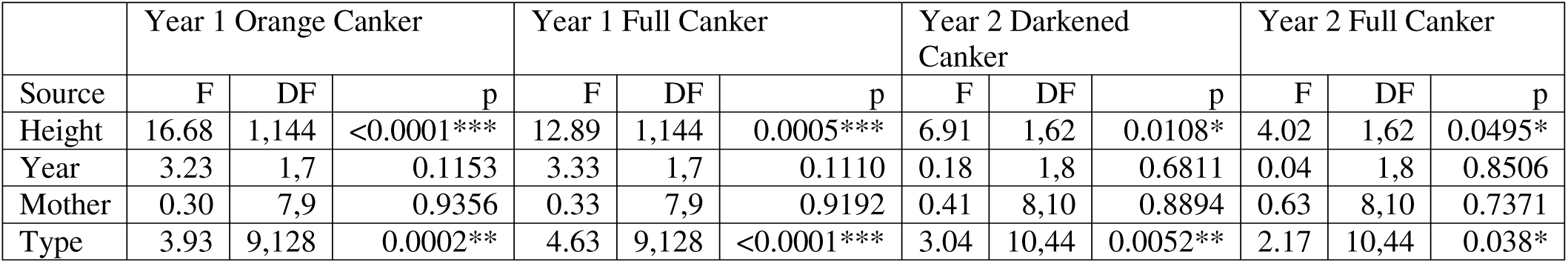
Combined full- and half-sib Nested Analysis of Covariance (ANCOVA) results evaluating the effects of Year, Mother Tree within Year, and Chestnut Type within Mother Tree (including Darling 54 and Wild-Type) on Orange and Full Canker length (Year 1 dataset) and Darkened and Full Canker length (Year 2 dataset). Due to the nested nature of the factors, ANCOVAs were run in stepwise fashion, first evaluating the effect of the covariate, height, on each branch inoculation assay via Simple Linear Regression analysis, and then running Nested ANOVAs on the residuals from the regressions. In order to fully meet model assumptions, Full Canker length measured in Year 1 and Height (both years) were square-root transformed, whereas Full Canker length measured in Year 2, Darkened Canker length and Orange Canker length were log-transformed.

**Supp.Table 3.** Combined full- and half-sib Nested Analysis of Covariance (ANCOVA) results evaluating the effects of Year, Pollen Source within Year, and Chestnut Type within Pollen Source (including Darling 54 and Wild-Type) on Orange and Full Canker length (Year 1 dataset) and Darkened and Full Canker length (Year 2 dataset). Due to the nested nature of the factors, ANCOVAs were run in stepwise fashion, first evaluating the effect of the covariate, height, on each branch inoculation assay via Simple Linear Regression analysis, and then running Nested ANOVAs on the residuals from the regressions. In order to fully meet model assumptions, Full Canker length measured in Year 1 and Height (both years) were square-root transformed, whereas Full Canker length measured in Year 2, Darkened Canker length and Orange Canker length were log-transformed.

|  | Year 1 Orange Canker |  |  | Year 1 Full Canker |  |  | Year 2 Darkened Canker |  |  | Year 2 Full Canker |  |  |
| --- | --- | --- | --- | --- | --- | --- | --- | --- | --- | --- | --- | --- |
| Source | F | DF | p | F | DF | p | F | DF | p | F | DF | p |
| Height | 16.22 | 1,134 | <0.0001*** | 13.56 | 1,134 | 0.0003*** | 6.91 | 1,62 | 0.0108* | 4.02 | 1,62 | 0.0495* |
| Year | 2.52 | 1,6 | 0.1634 | 1.54 | 1,6 | 0.2611 | 0.39 | 1,6 | 0.5563 | 0.07 | 1,6 | 0.8059 |
| Pollen | 0.38 | 6,6 | 0.8653 | 0.69 | 6,6 | 0.6704 | 0.18 | 6,8 | 0.9752 | 0.30 | 6,8 | 0.9197 |
| Type | 5.31 | 6,122 | 0.0001*** | 6.64 | 6,122 | <0.0001*** | 2.89 | 8,48 | 0.0103* | 2.46 | 8,48 | 0.0256* |

**Supp.Table 4.**
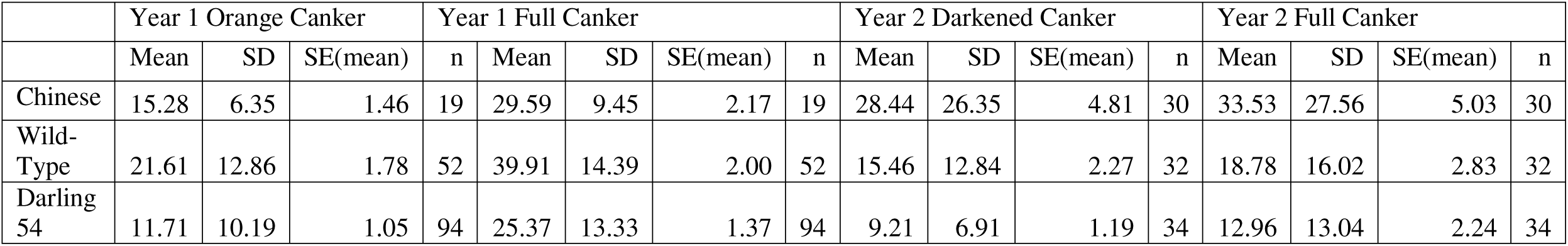
Summary inoculation assay statistics for Chestnut Types measured in Years 1 and 2.

## Footnotes

7 The research team at State University of New York Environmental Science and Forestry College that produced the genetically engineered American chestnut was headed by the late Dr. William Powell and now by Dr. Andrew Newhouse.

8 Animal and Plant Health Inspection Service, US Department of Agriculture

9 In 2020, leaf samples from three mother trees in Maine that phenotypically appeared to be American chestnut were part of a larger group of samples sent to genomics labs at the University of Notre Dame and Virginia Tech University for chestnut species analysis. The independent results from the two labs closely matched, revealing that one of the mother trees was approximately 21% European chestnut (*Castanea sativa* Mill.) while two others were 40% European chestnut. When bred with 100% American chestnut pollen, offspring from these three trees that are included in this study are on average 10.5-20% European chestnut.

